# Vision affects gait speed but not patterns of muscle activation during inclined walking – a virtual reality study

**DOI:** 10.1101/2020.07.28.226118

**Authors:** Amit Benady, Sean Zadik, Oran Ben-Gal, Desiderio Cano-Porras, Atalia Wenkert, Sharon Gilaie-Dotan, Meir Plotnik

**Affiliations:** Center of Advanced Technologies in Rehabilitation Sheba Medical Center, Israel; St George’s University of London medical school, Sheba Medical Center, Israel; School of Optometry and Vision Science, Bar Ilan University, Ramat Gan, Israel; Brightlands Institute for Smart Society - BISS, Maastricht University, Maastricht, The Netherlands; Dept. of Physiology and Pharmacology, Sackler School of Medicine, Tel Aviv University, Tel Aviv, Israel; UCL Institute of Cognitive Neuroscience, London, UK; Sagol School of Neuroscience, Tel Aviv University, Tel Aviv, Israel

**Keywords:** Locomotion, inclined surfaces, visuomotor integration, electromyography, internal model of gravity, visual dependency, virtual reality

## Abstract

While walking, our locomotion is affected by and adapts to the environment based on vision-based and body-based (vestibular and proprioception) cues, all contributing to an “Internal Model of Gravity”. During surface inclination transitions, we modulate gait to counteract gravitational forces by braking during downhill walking to avoid uncontrolled acceleration or by exerting effort to avoid deceleration while walking uphill. In this study, we investigated the role of vision in gait modulation during surface inclination transitions by using an immersive large-scale Virtual Reality (VR) system equipped with a self-paced treadmill and projected visual scenes that allowed us to modulate physical-visual inclinations congruence parametrically. Gait speed and leg muscle electromyography (EMG) were measured in 12 healthy young adults. In addition, the magnitude of subjective visual misperception of verticality was measured by the rod and frame test. During virtual (non-veridical) inclination transitions, vision modulated gait speed after transitions by (i) slowing down to counteract the excepted gravitational ‘boost’ in virtual downhill inclinations and by (ii) speeding up to counteract the expected gravity resistance in virtual uphill inclinations. These gait speed modulations were reflected in muscle activation intensity changes and associated with subjective visual verticality misperception. However, temporal patterns of muscle activation, which are significantly affected by real gravitational inclination transitions, were not affected by virtual (visual) inclination transitions. Our results delineate the contribution of vision to functional locomotion on uneven surfaces and may lead to enhanced rehabilitation strategies for neurological disorders affecting movement.

**Significance statement:** A crucial component of successful locomotion is maintaining balance and speed while walking on uneven surfaces. In order to reach successful locomotion, an individual must utilize multisensory integration of visual, gravitational, and proprioception cues. The contribution of vision to this process is still unclear, thus we used a fully immersive virtual reality treadmill setup allowing us to manipulate visual (virtual) and gravitational (real) surface inclinations independently during locomotion of healthy adults. While vision modulated gait speed for a short period after inclination transitions and this was predictive of individual’s visual dependency, muscle activation patterns were only affected by gravitational surface inclinations, not by vision. Understanding the vision’s contribution to successful locomotion may lead to improved rehabilitation for movement disorders.

## Introduction

In order to maintain stability during bipedal locomotion, the muscle system continuously adapts to changing walking conditions via a complex motor process that relays on multisensory integration (Wall-Scheffler CM, Chumanov E and B, 2011) that involves whole body adaptations(Kimel-Naor et al., 2017; Cano Porras et al., 2020). Particularly, walking on inclined surfaces lead to significantly different patterns and magnitudes of activation in lower limb muscles, as compared to leveled walking. This is evident by changes in patterns and magnitude of increased (e.g., rectus femoris, tibialis anterior) or decreased (e.g., gastrocnemius muscle) activation of specific muscles during downhill walking (Lay et al., 2007; Pickle et al., 2016) or by increased muscles activation (e.g., gastrocnemius, hamstring and rectus femoris) during uphill walking (Pickle et al., 2016; Janshen et al., 2017).

Inclined walking involves adjustment to gravitational forces. The *exertion effect* (uphill walking) and the *braking effect* (downhill walking) have been previously described and quantified (Cano Porras et al., 2020). Briefly, during uphill walking, the exertion effect counteracts the gravitational deceleration and allows the walker to maintain roughly stable gait speed pace, which is lower than *natural* speed during leveled walking (Sun et al., 1996; Sinitski et al., 2015; Kimel-Naor et al., 2017). During downhill walking, the braking effect prevents (gravitational-driven) uncontrolled speeding-up and allows the walker to descend in a steady gait speed, either faster or slower than walking on a leveled surface(Sun et al., 1996; McIntosh et al., 2006; Kimel-Naor et al., 2017; Cano Porras et al., 2020).

It is suggested that locomotor modulation following perceived gravitational changes while walking is mediated through an ‘*internal model of gravity*’(Merfeld et al., 1999; Campos et al., 2014; Lacquaniti et al., 2014; Balestrucci et al., 2017) that integrates multisensory available cues such as vestibular, proprioceptive (aka body-based cues) and visual cues (Figure 1). The exact role of vision in this process is yet unclear. To evaluate the relative contribution (‘weight’) of vision to the internal model of gravity during locomotion there is a need to manipulate visual cues independently of body-based cues. In a recent study, we found that vision modulates gait speed, postural adjustment and spatio-temporal gait parameters shortly after transition to a virtual inclination for approximately 20s. Yet it remains unclear whether individuals which are more dependent on visual inputs than others, would respond by greater modulations in gait speed following a virtual visual inclination. Furthermore, it is unclear whether patterns of muscle activation are also affected by mere virtual inclinations.

**Figure 1.**
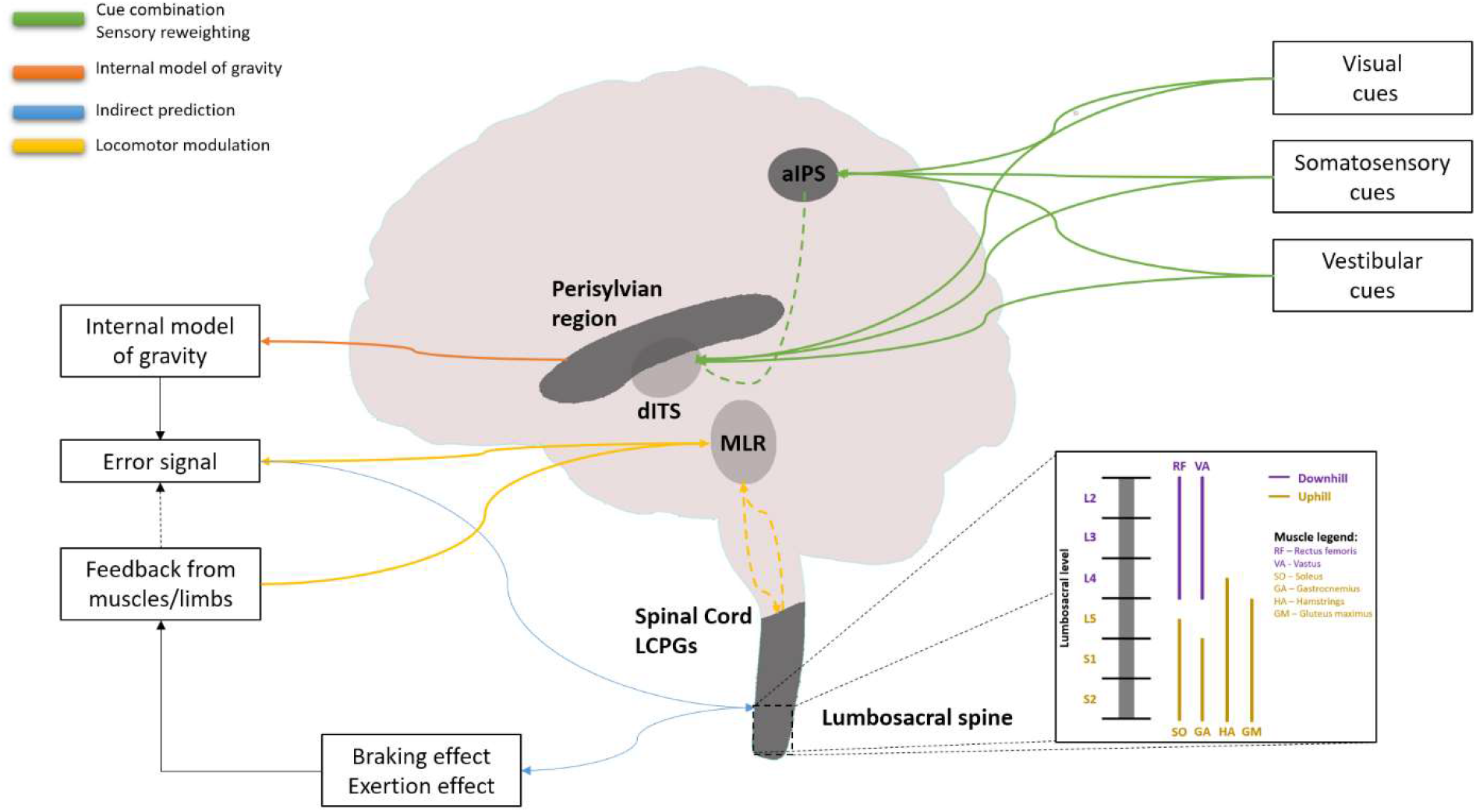
A neural model that putatively accounts for locomotor modulations following perceived gravitational changes. This predictive system relies on the internal model of gravity, which is assumed to regulate locomotion according to physical laws of gravity. Indirect prediction may occur through spinal reflexes, which optimize energy consumption, or through central pattern generators (Pearson, 2004; Snaterse et al., 2011). Pattern generators are distributed in the spinal cord with specific segments corresponding to specific muscle innervations (Ivanenko et al., 2006). The synergic interactions among locomotion central pattern generators (LCPGs) are highly robust, time-adaptive and adjust to changing environmental contexts (Choi and Bastian, 2007). LCPGs control the activation of lumbosacral alpha motoneurons responsible for the activation of muscles with specialized functional roles in inclined walking (Ivanenko et al., 2006; Flash and Bizzi, 2016; Pickle et al., 2016). Therefore, LCPGs may be accountable for the synchronized muscle activations that facilitate braking and exertion effects. The motor cortex, brainstem and cerebellum all send inputs to the locomotor region (LMR) in the midbrain through the LCPGs. The LMR receives afferent proprioceptive feedback from the limbs and muscles (Prochazka and Ellaway, 2012). This feedback loop may represent one key component for gradually correcting the error arising from incongruent sensory inputs to the IMG. Relevant brain regions are shown. Right lower panel shows a schematic representation of the lumbosacral maps of LCPGs involved in the activation of muscles that play critical roles during inclined walking (Ivanenko et al., 2006; Pickle et al., 2016). The Green lines demonstrate brain areas involved in sensory integration and reweighting, the orange line demonstrate a region proposed to subserve the IMG. Blue lines demonstrate pathways involved in indirect prediction, and gold lines indicate proposed mechanisms of locomotor regulation and action. Dashed lines represent connectivity between areas in the central nervous system (CNS). MLR: mesencephalic locomotor region; LCPGs: locomotion central pattern generators, dITS: dorsal part of the inferior temporal sulcus [homologous to macaque dorsal medial superior temporal area], aIPS: anterior part of the intraparietal sulcus [homologous to macaque ventral intraparietal area (Grefkes and Fink, 2005)]. Adapted from Cano Porras et al., (Cano Porras 2020).

Visual field dependence in the context of locomotion is considered as the level of reliance on visual cues in comparison to body-based cues (Isableu et al., 1998; Willey and Jackson, 2014). A common method to assess visual field dependency is through the rod and frame test which is assumed to estimate the extent of subjective misperception of visual verticality (Lopez et al., 2006; Isableu et al., 2008; Bagust, 2013). Individual differences in visual field dependence have been reported (Anon, n.d.; Kaleff et al., 2011) and it has been suggested to relate to balance in patients and populations with balance related disorders(Lord and Webster, 1990; Bonan et al., 2006, 2007; Crevits et al., 2007).

In the present study we aimed to explore whether the gait speed modulations triggered by mere visual cues after transitioning to virtual inclined surface walking are accompanied by changes in muscle activation patterns typical to those triggered by veridical (gravitational) surface inclination transitions. Predictions based on our previously proposed putative neural model support this hypothesis (Cano Porras et al., 2020) (Figure 1). Furthermore, we investigated whether the level of subjective visual verticality misperception was associated with the change in gait speed following virtual (visual) surface inclination change.

## Materials and Methods

### Participants

Twelve young, healthy adults (mean age ± SD: 26.53±3.09 years old, 6 males) participated in this study. Exclusion criteria were cognitive limitations, physical and visual restrictions, and any sensorimotor impairments that could potentially affect locomotion or the ability to adhere to instructions. The Institutional Review Board for Ethics in Human Studies at the Sheba Medical Center, Israel, approved the experimental protocol, and all participants signed a written informed consent prior to entering the study.

### Apparatus

#### Virtual reality system

Experiments were conducted in a fully immersive virtual reality system (CAREN High End, Motek Medical, The Netherlands) containing a moveable platform with six degrees of freedom (Kimel-Naor et al., 2017). A treadmill in a self-paced mode was embedded in the moveable platform allowed participants to adjust it according to their preferred gait speed (Plotnik et al., 2015). A motion capture system (Vicon, Oxford, UK) tracked the three-dimensional coordinates of 41 passive markers located on each participant’s body with a sampling frequency of 120 Hz and spatial accuracy of 1 mm.

#### Physiological measures recording device – Electromyography (EMG)

Electrical activity was recorded at 1024 Hz (eego^tm^ sports, ANT neuro, the Netherlands) from four right lower limb muscles: Tibialis Anterior (TAR) Gastrocnemius (GCR), Rectus Femoris (RFR) and Hamstring lateralis (HLR), as these muscles have been previously described to alter their activation pattern during uphill and/or downhill walking. The location of electrodes followed the SENIAM guidelines (Hermens et al., 1999). Pairs of self-adhesive bipolar electrodes were attached over the muscle bellies with 2 cm between electrodes. Prior to recording, a calibration stage was conducted by asking participants to precisely move the relevant muscle for confirming reliable activation patterns on the control monitor.

#### VR version of the Rod and Frame Test

The rod-and-frame test estimates subjective visual verticality misperception. Specifically, the test measures how visual perception of the orientation of a central bar (rod) is influenced by the orientation of a peripheral visual reference frame around it. Implementation of the test was conducted in our lab using Unity software and C# scripting. The participants sat upright wearing head mount device (HMD) VR glasses (HTC VIVE, HTC; New Taipei City, Taiwan) and were told not to move or tilt their head during the test. The VR environment consisted of a white frame at a specific orientation and a white rod inside it with its own orientation; both presented on a black background. A sequence of 28 trials was presented during which the frame was initially at one of seven possible random positions: 0/±10/±20/±30 degrees (0 was vertical, + was clockwise). Each of these initial frame positions was presented four times (Bagust, 2005). In addition, the initial angle of the rod was randomized (sampled from 0-180 degrees range distribution). The participants aimed to orient the rod perpendicular (i.e., vertical) to the true horizon, regardless of the surrounding frame’s orientation. This was achieved by rotating the rod around its center in a clockwise or counterclockwise direction using the VR system’s remote control. It is critical to note that the surrounding frame was unchanged by this manipulation. Once the participants perceived it as being vertical, they responded by pressing a button on the remote control, which leads to the clearing of the display and the beginning of another trial immediately.

### Procedure

#### VR rod-and-frame test

After filling the informed consent, the first part of the experiment was to perform the rod and frame test. First the team began by assuring that the participant felt comfortable with the head mount device (HMD). Next, the participant underwent a short practice trial to confirm that he/she fully understood the task. Following the practice trial, 28 test trials were conducted and measured accurately. It is critical to note that the tests was not limited in time and typically lasted 10 minutes, including the practice trial.

#### Gait trials in large-scale VR system

##### Habituation period to walking in self-paced mode during leveled and inclined surfaces

A safety harness secured the participant to a metal frame on the moveable platform (Figure 2). The first part of the habituation was to familiarize the participant with the self-paced mode of the treadmill, which involved 10-15 minutes of leveled walking, aiming to provide practice of increasing and decreasing speed until he/she mastered the walking. The second part of the habituation involved connecting the EMG electrodes and calibrating the motion capture system. The third and final part of the habituation included one walking trial of the three possible inclinations (i.e., leveled, uphill, and downhill walking) when the visual and the gravitational cues were synchronized (‘congruent’ conditions; see more details below). Each trial lasted three to four minutes.

**Figure 2.**
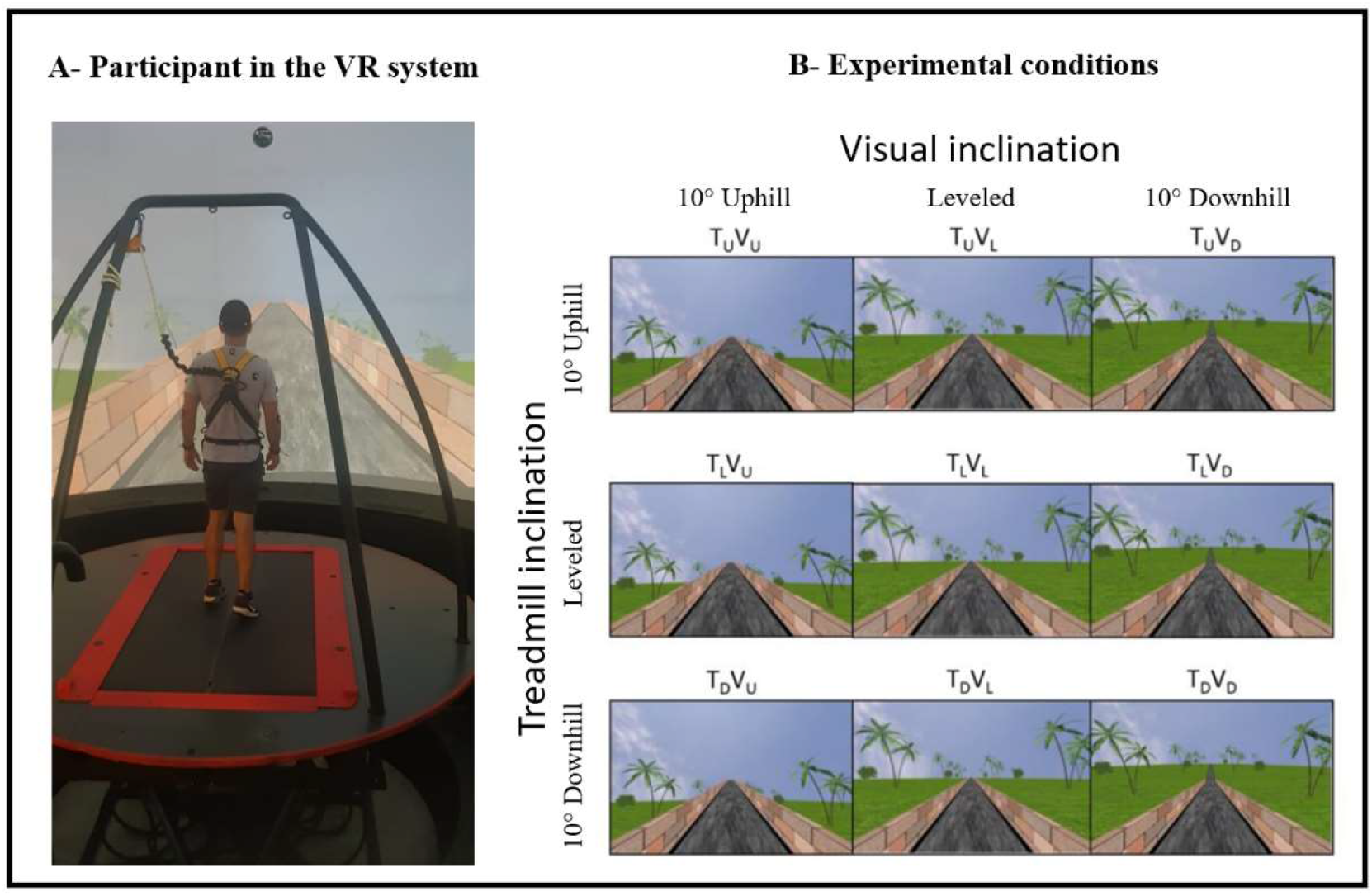
Apparatus (A) and the main experimental conditions (B). **(A)** A fully immersive virtual reality system containing an embedded treadmill synchronized with projected visual scenes, wherein this example, the treadmill is leveled, and the vision is uphill (see T_L_V_U_ in B). **(B)** Main experimental conditions. Following leveled walking and after reaching steady-state velocity (SSV) and maintaining it for 12 seconds, a transition (5s) occurred to one of eleven different conditions presented in random order, lasting 65 seconds, in which the inclination of the treadmill and/or visual scenes (V) transitioned to 10° uphill (_U_), remained leveled (_L_) or transitioned to −10° downhill (_D_). Rows represent treadmill inclination changes, and columns represent visual scene inclination changes. Visual scene inclination effect was achieved by the road appearing above (uphill), below (downhill), or converging (leveled) with the line of the horizon. In addition, the peripheral greenery is exposed more (downhill) or less (uphill) by the road. Vision-treadmill congruent conditions appear on the diagonal (T_L_V_L_ for continued leveled walking, T_U_V_U_ and T_D_V_D_ for uphill and downhill walking, respectively). Vision-treadmill incongruent conditions below the diagonal represent conditions with visual scene inclination more positive than the treadmill’s (T_L_V_U_, T_D_V_U_, T_D_V_L_), and above the diagonal visual scene inclination more negative than the treadmill’s (T_U_V_L_, T_U_V_D_, T_L_V_D_). See Methods for more details.

##### Gait experiments

The participants were informed that they would perform several short gait trials with short intervals between them. They were instructed to walk “as naturally as possible” and that “inclinations may occur during walking”. It is important to note that the condition did not affect the beginning stage of the trials, as every participant was explicitly instructed to begin from a standstill position and then progress into walking with both the treadmill and the visual scene leveled until reaching steady-state velocity. Once reaching steady-state velocity, a 5s long transition of the treadmill and/or visual scene occurred (except in the congruent level condition, where no transition occurred). Post transition, participants walked for additional 65s until the treadmill slowed down and stopped altogether. By convention, we refer to the transition start time as time zero (t=0).

##### Experimental conditions

The protocol included eleven experimental conditions that the participant encountered in random order. Inclination of the treadmill (T) and/or visual scenes (V) transitioned to 10° uphill (U), remained leveled at 0° (L) or transitioned to −10° downhill (D). Figure 2 depicts the main 3×3 experimental design conditions where rows represent treadmill (T) inclination and columns represent visual scene (V) inclination (Note that inclination of 10° is considered steep (Proffitt et al., 1995). Treadmill and visual scene congruent conditions that served as baseline appear on the diagonal for leveled (T_L_V_L_), uphill (T_U_V_U_), and downhill (T_D_V_D_) walking. Visual scene-treadmill incongruent conditions include the following visual scene manipulations: for treadmill uphill inclination, the vision was leveled (T_U_V_L_) or downhill (T_U_V_D_), for during treadmill leveled inclination, the vision was uphill (T_L_V_U_) or downhill (T_L_V_D_), and lastly, for treadmill downhill inclination, the vision was leveled (T_D_V_L_) or uphill (T_D_V_U_).

Since we hypothesized finding exertion and braking effects during virtual (visual) inclination transitions, which could possibly affect muscle activation patterns, there were two additional control conditions without treadmill or visual scene transitions. Participants were asked to increase (i.e., T_L_V_L_-increase) or decrease (i.e., T_L_V_L_-decrease) their speed voluntarily until the experimenter (A.B.) instructed them to maintain their current speed until the end of the trial. The target speeds were an increase of ~15% and a decrease of ~20% from individuals’ steady-state velocity (SSV), and the experimenter monitored this via the control computer. The quantitative values of the changing speeds were chosen to match the gait speed increase (in T_L_V_U_ condition) and the gait speed decrease (in T_L_V_D_ condition) observed in our previous study (Cano Porras et al., 2020). The purpose of these two conditions was to control the possibility that muscle activity pattern changes under virtual inclinations would be due to gait speed changes rather than to the change of gravitational forces.

### Outcome measures

#### Gait speed related variables

To assess the post-transition effects on gait speed, we followed the methodologies introduced by Cano-Porras et al. (Cano Porras et al., 2020). We looked at (i) the magnitude of the peak/trough of gait speed relative to the steady-state velocity (SSV; presented in %); and (ii) the time of this peak from the start of transition (seconds). For a full derivation of these parameters, see *Supplementary Methods*.

#### Standardized response to virtual inclination

To compute this metric, we used data from the incongruent T_L_V_U_, T_L_V_D_, T_D_V_U_, T_U_V_D_ conditions. The absolute values of the maximal relative (i.e., percent) change were calculated for each participant with respect to the SSV.

#### Subjective verticality misperception index

For each trial, the degree of deviation of the rod from the true vertical was measured and recorded as the position error. For each participant, the mean position error for 7 different frame angles were calculated. Data from all participants were grouped by the frame angle (Bagust, 2005). We defined the rod and frame index as the average angle of deviation of the rod from the true vertical when the frame was projected at ±20 degrees (8 trials in total, 4 trials of +20° and 4 trials of −20°). This parameter allowed us to evaluate individual differences in gravitational misperception.

#### EMG patterns evaluation

EMG traces were parsed in the temporal domain to gait cycles based on heel strike detection. Briefly, EMG traces from several gait cycles from the pre-transition period were averaged for each participant for each condition. Similarly, this was done for the post-transition period. In order to compare the condition effect, two specific parameters were studied:

1. Magnitude – the summation (i.e., area under the curve) of the EMG traces.
2. Similarity of EMG activation pattern – For each participant and condition, the post-transition EMG trace was correlated with the pre-transition EMG trace (Pearson correlation). A high correlation value indicates a small change in the pattern of activation (see *Supplementary Material* for more details.

### Statistical analyses

In order to test the differences between the walking conditions, we analyzed each dependent variable separately. To analyze the magnitude of activation, we compared the mean of each variable with each muscle pre- and post-transition using a two-tailed paired T-test. To analyze the similarity of pattern, we compared the correlation of each muscle pre- and post-transition. A p-value of less than or equal to 0.05 was considered statistically significant. Pearson correlation coefficient was computed to evaluate the relationship between subjective verticality misperception index and the standardized response to virtual inclination. Data were extracted using MATLAB software (MATLAB R2017a script (The MathWorks, Inc, Natick, MA) and analyzed using SPSS software (v25, IBM).

## Results

### The effects of visual cues on walking in physical and virtual inclinations

We first sought to replicate earlier findings (Cano Porras et al., 2020) of how vision behaviorally influences gait speed for a short period after a visual surface inclination transition occurs, regardless of the veridical inclination (see demonstration video in *Supplementary Material*). Figure 3 depicts the average normalized self-paced gait speed relative to steady-state velocity for each condition, one-minute pre, and one-minute post inclination transition. Briefly, in the congruent transition conditions (T_U_V_U_ and T_D_V_D_) there was a gradual monotonic change for downhill and a steep monotonic change for uphill walking. In contrast, the incongruent conditions were characterized by peaks (or troughs) in gait speed between 5-15s after the virtual transition, which was determined by the vision-treadmill discrepancy direction (troughs when virtual inclination was lower than treadmill’s inclination, positive peak when virtual inclination was higher than treadmill’s inclination). It is critical to note that the vision-treadmill discrepancy direction was proportional to the vision-treadmill discrepancy magnitude. This explains why a 20° discrepancy leads to a bigger peak than a 10° discrepancy.

**Figure 3.**
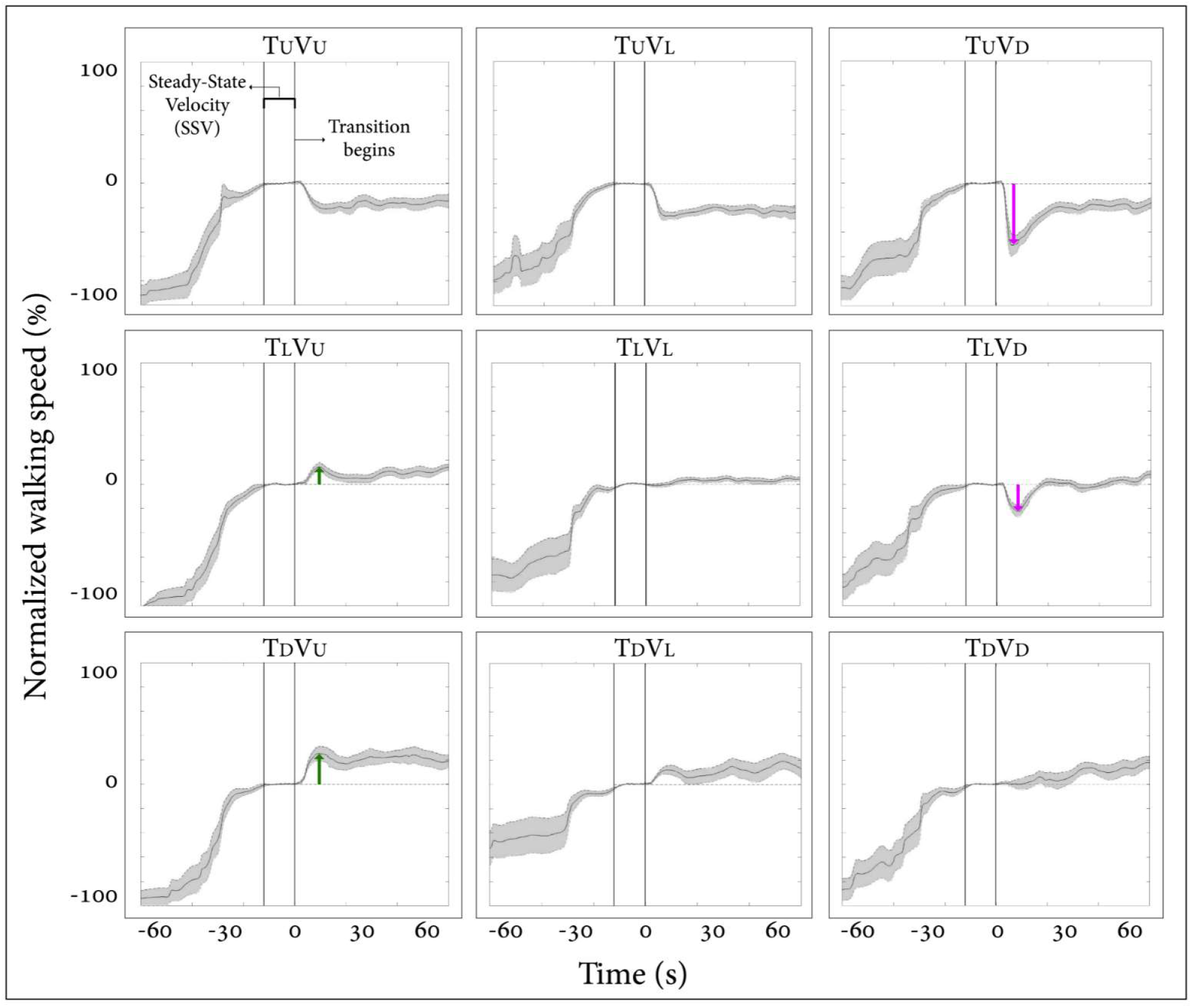
Post-transition adaptation of gait speed. Average self-paced gait speed (N=12) relative to steady-state velocity (in %) for each condition. Time zero demarcates the end of the steady-state velocity period, after which a 5s transition of the treadmill and/or visual scene occurred. Gray shading represents the standard error. Green arrows represent the peak of excretion effects in visual uphill walking, and red arrows represent the peak of braking effects in visual downhill walking. Irrespective of treadmill inclination upon transition, there is a tendency for a virtual (visual) downward transition to be followed by a decrease in gait speed and, to a lesser extent, for an upward visual transition to be followed by an increase in gait speed. Conditions: treadmill (T) and/or visual scenes (V) inclination transitioned to 10° uphill (U), remained leveled at 0° (L), or transitioned to −10° downhill (D).

We examined how vision affected gait speed by examining how parametric visual inclination changes affected gait speed in conditions where the treadmill inclination remained constant (see rows in Figure 3, (where each row represents a fixed treadmill inclination change). For the T_U_ conditions (when treadmill transition changed to 10° uphill, upper row of Figure 3), there was a gradual decrease in gait speed during the first 20 seconds after transition (cf. left T_U_V_U_ panel with smallest decrease, middle T_U_V_L_ panel with a medium decrease, and right T_U_V_D_ panel with largest decrease after which the gait speed returned roughly to gravitational-based gait speed regardless of the visual (virtual) inclination (see values and comparison to Cano Porras et al. in the Supplementary Material). Similarly, for the T_L_ conditions (when the treadmill remained leveled at 0°, middle row of Figure 3), there was a gradual change in gait speed in the first 20 seconds after the virtual transition (cf. left T_L_V_U_ panel with an increase (visually induced exertion effect), middle T_L_V_L_ panel with no change, and right T_L_V_D_ panel with a decrease (visually induced braking effect)) after which the gait speed returned roughly to gravitational-based gait speed regardless of the visual (virtual) inclination. For the T_D_ conditions (when treadmill transition changed to −10° downhill, lower row of Figure 3), the same pattern of a gradual change in gait speed in the first 20 seconds after the virtual transition was observed (cf. left T_D_V_U_ panel with an increase, middle T_D_V_L_ panel with a smaller increase and right T_D_V_D_ panel with no change). Following that, gait speed returned roughly to the gravitational-based level regardless of visual inclination. In summary, this analysis indicates that visual input in the form of virtual inclinations affected and modulated gait speed in a parametric fashion of 5-20s after the transition occurred when the treadmill transition was kept constant.

### Relation between visual modulation of gait speed during visual-physical incongruent conditions and subjective misperception of verticality

We hypothesized that the magnitude of visual modulation on gait speed during virtual surface inclination changes varies across people and may be related to an individual’s subjective visual misperception of verticality. Therefore, for each participant we calculated the magnitude of visual modulation on gait speed based on the gait speed changes in the incongruent conditions (i.e., standardized response to virtual inclination, see Methods for calculation). An additional factor for our calculation was the index of subjective visual misperception of verticality estimated by the rod and frame illusion magnitude (see Methods). We found a significant correlation when comparing these two independently obtained measurements (Figure 4; Pearson’s R=0.76, p= 0.004 (t(10)= 3.69)), suggesting that they may rely on associated mechanisms.

**Figure 4.**
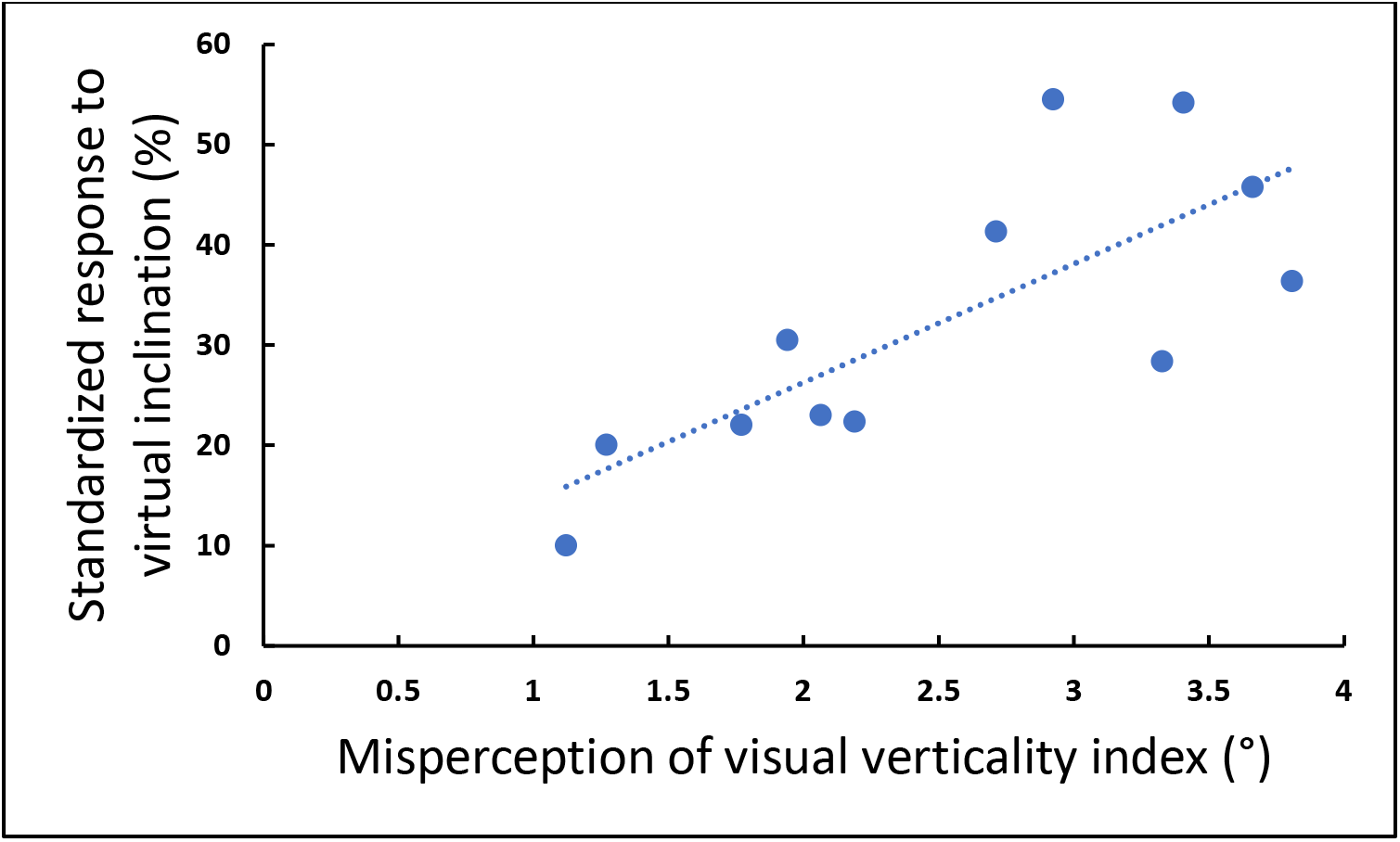
Visual field dependence relative to misperception of visual verticality. The extent of visual field dependence during locomotion with inclined surfaces is linked to the misperception of visual verticality. The X-axis represents the misperception of the visual verticality index assessed by the psychophysical rod and frame test while seated. The Y-axis represents the standardized response to virtual inclination based on incongruent conditions, and each circle represents a participant (see Methods). A significant correlation was found between these measures (Pearson’s R=0.76, p=0.004 (t(10)=4.075), suggesting that they may be relying on joint mechanisms. The dotted line represents the linear regression line Y= 0.472X + 0.1.

### The effects of visual and physical transitions on muscle activation patterns

Consistently with our primary objective, and to test our central hypothesis, we examined whether visual cues affect muscle activation patterns (besides modulating behavioral gait measures). To that end, we analyzed pre- and post-transition EMG muscle activation patterns of four different muscles (Tibialis Anterior Right (TAR), Gastrocnemius Right (GCR), Rectus femoris Right (RFR) and Hamstring Lateral Right (HLR)). The reasoning for choosing those specific muscles was based on their activation patterns, which were found to be modulated by gravitational surface inclination transitions (Lay et al., 2007; Pickle et al., 2016).

As can be appreciated from the top row of Figure 5, each of the four muscles presented a unique activation pattern during the leveled walking stage. Also, as expected for the leveled walking (control condition), the pre-(leveled in blue) and post-transition (leveled in orange), traces were comparable. Furthermore, in the treadmill-vision congruent transitions (uphill T_U_V_U_ in the second row, and downhill T_D_V_D_ in the third row, Figure 5) there were significant changes between the pre-transition (in blue) muscle activation patterns (note that it is comparable to that of T_L_ conditions in the upper row) and the post-transition muscle activation patterns (in orange). It is critical to note that the significant changes took place in all four muscles, except HLR in the T_D_V_D_ condition (see Table 1 for p-values).

**Figure 5.**
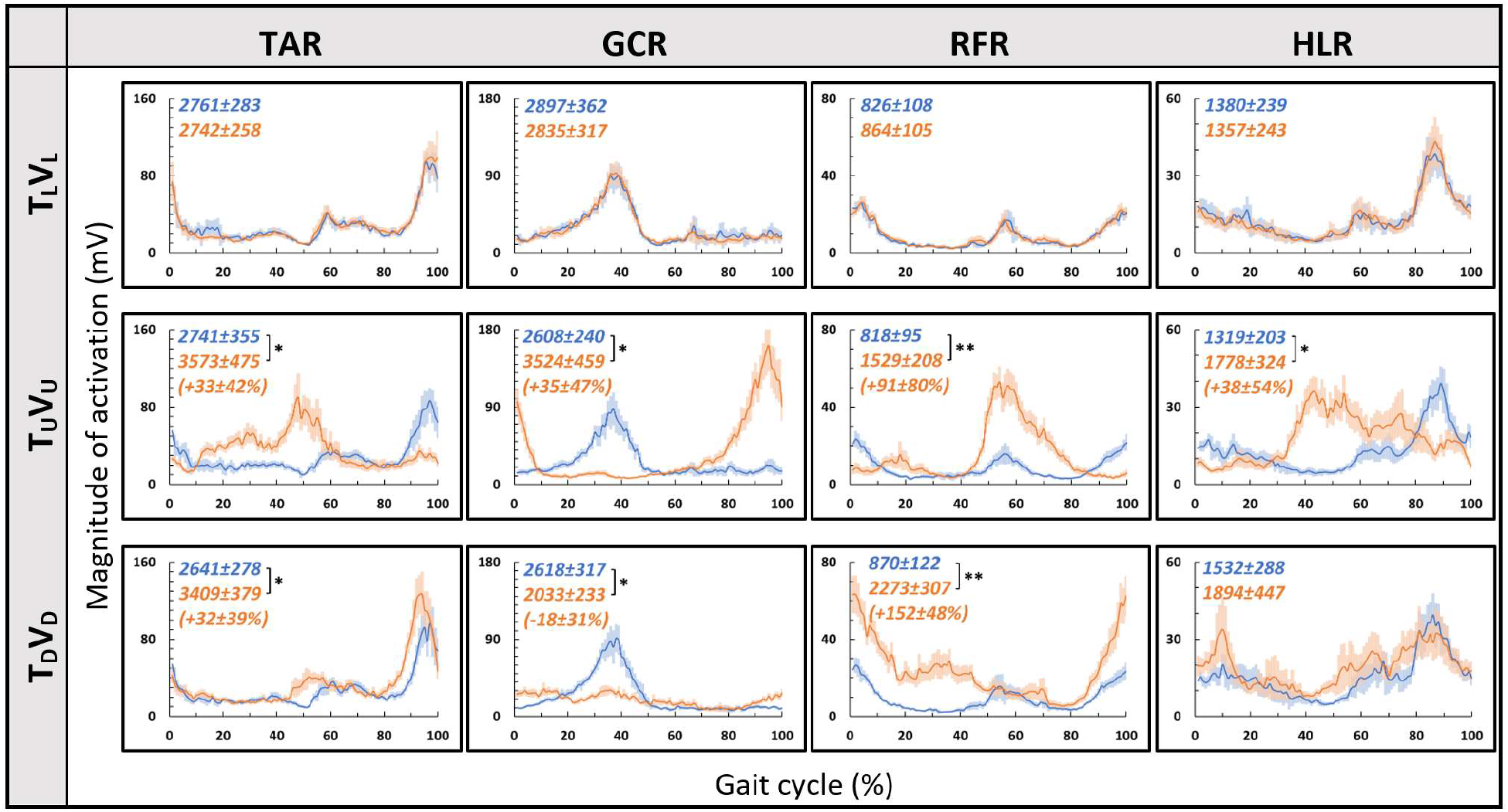
Pre- vs. post- EMG muscle activation patterns for visual-physical congruent transitions. Grand averages (across participants, N=12) of muscle activation patterns (Y-axis, in mV) plotted against a time-normalized gait cycle on the X-axis (in % of the gait cycle) in blue (shaded areas represent SE) for pre-transition and in orange (shaded areas represent SE) for 20 seconds post-transition. Numerical values at the top left of each panel represent averaged summation (area under the curve) magnitude of activation ± SE for pre-transition (blue) and post-transition (orange), asterisks denote a significant post- vs. pre-transition change (two-tailed, paired T-test, P<0.05=*, P<0.01=**) which appears below then (in %) in parentheses. Upper row displays pre- and post-transition leveled walking for each of the four muscles, and as expected, pre vs. post are comparable for each of the muscles. Note that in both uphill (middle row) and downhill (bottom row) transitions, the pre-transition (blue curve) is comparable to the leveled condition (blue in the upper row). Post-transition significant changes are evident for inclined transitions (uphill or downhill) for all four muscles apart from the HLR downhill transition (bottom right). TAR= Tibialis Anterior Right, GCR= Gastrocnemius Right, RFR= Rectus Femoris Right. HLR= Hamstring Lateralis Right. See Results and Methods for more details.

**Table 1.** The grand averaged (N=12) p-value for summation magnitude of muscle activation between pre- and post-transition for all experimental conditions. Each row represents one walking condition and each column represents a specific muscle. A significant value indicates that there was a significant change in the summation of magnitude of muscle activation following the transition of the treadmill and/or visual scene. P value < 0.05=*, P value <0.01=**.

|  | **TAR** | **GCR** | **RFR** | **HLR** |
| --- | --- | --- | --- | --- |
| **T<sub>L</sub>V<sub>L</sub>** | 0.868 | 0.579 | 0.290 | 0.669 |
| **T<sub>L</sub>V<sub>U</sub>** | **0.005** | **0.001** | **0.01** | **0.01** |
| **T<sub>L</sub>V<sub>L</sub> Increase** | **0.0017** | **0.000** | **0.000** | **0.000** |
| **T<sub>L</sub>V<sub>D</sub>** | 0.110 | **0.004** | 0.978 | 0.203 |
| **T<sub>L</sub>V<sub>L</sub> Decrease** | 0.060 | **0.038** | 0.858 | **0.024** |
| **T<sub>U</sub>V<sub>U</sub>** | **0.021** | **0.013** | **0.001** | **0.004** |
| **T<sub>U</sub>V<sub>D</sub>** | **0.009** | 0.064 | **0.000** | **0.023** |
| **T<sub>D</sub>V<sub>D</sub>** | **0.015** | **0.033** | **0.000** | 0.173 |
| **T<sub>D</sub>V<sub>U</sub>** | **0.011** | 0.087 | **0.000** | **0.048** |

We then examined whether parametric changes of the visual cues (i.e., the visual (virtual) inclination) would affect muscle activation patterns for a specific treadmill transition inclination. As illustrated in Figure 6, for the leveled treadmill (T_L_, at 0°), the parametric manipulation of visual cues did not significantly affect the activation patterns of any of the four muscles as apparent by the similarities between the orange (post-transition) and blue (pre-transition) traces. These qualitatively detected similarities were confirmed by a series of correlation analyses (Table 2). As for muscle activation magnitude, it can be seen that during virtual uphill walking, all four muscles exhibited increased activation during the post-transition period compared to the pre-transition period. Addressing the virtual downhill walking, only the GCR exhibited a significant decrease in activation in the post-transition period compared to the pre-transition period.

**Figure 6.**
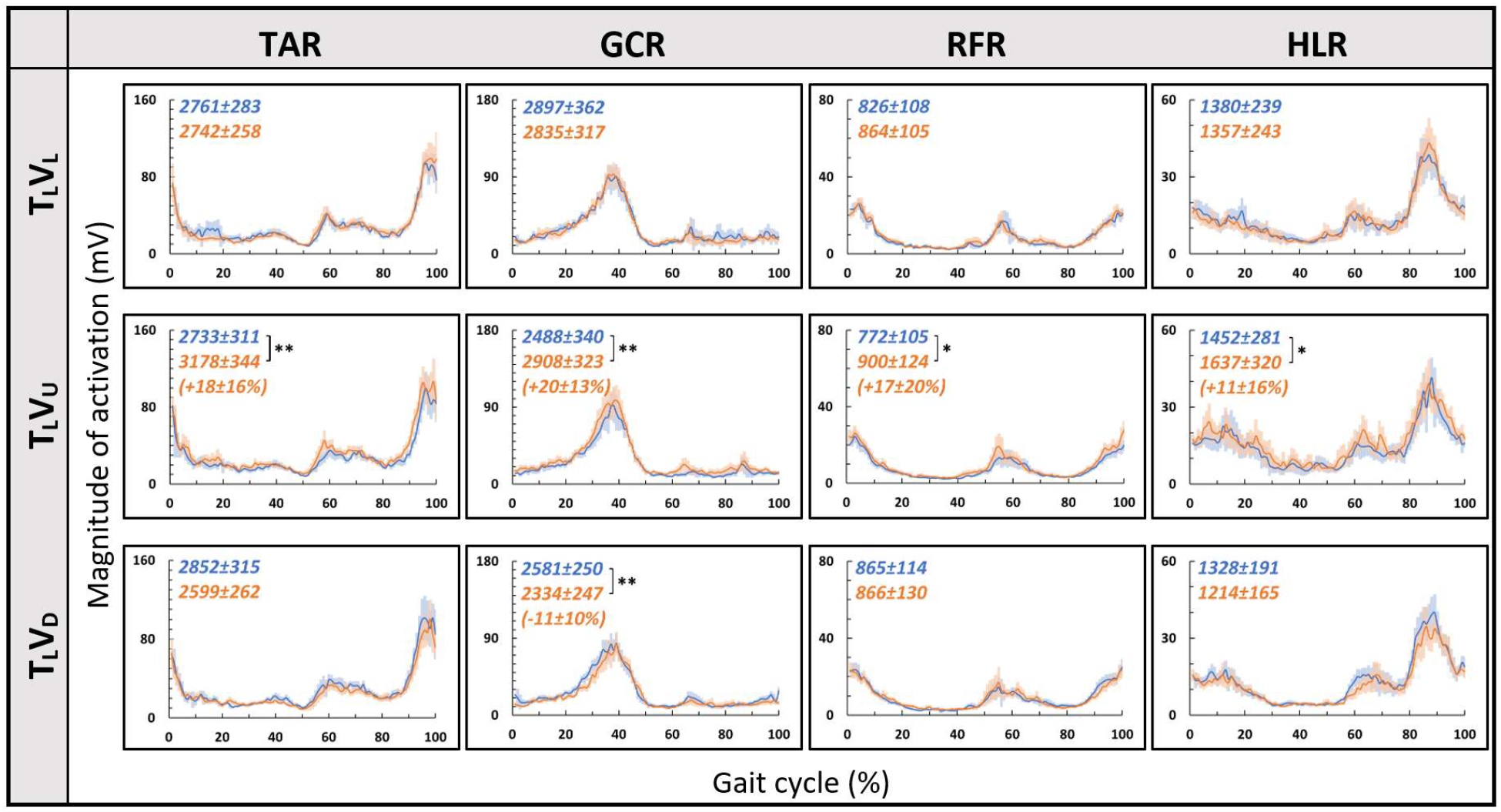
Visual inclinations do not affect muscle activation patterns for leveled treadmill inclination. X-axis represents the percentage of the gait cycle (%), and the Y-axis represents the magnitude of muscle activation (mV). Top row-leveled vision (0°, as in Figure 5 top row), middle row-vision up +10°, bottom row-vision down −10°. The blue lines represent the pre-transition and the orange lines represent post-transition activity. Numerical values superimposed on the panels represent the average magnitude of activation ± SE for pre-transition (blue) and post-transition (orange), and in parentheses significant changes (in %) between these conditions. T-test, P<0.05=*, P<0.01=**, (N=12). Despite the significant effects in the magnitude, we found that the EMG temporal patterns were highly comparable, see correlation analyses in Table 2.

**Table 2.** Correlations between pre- and post-transition. Average Pearson correlations (±SE) for each experimental condition. High correlations demonstrate comparable (pre and post transition) temporal patterns of muscle activation. All walking conditions show similar correlations according to the treadmill inclination, regardless of the visual scene inclination, suggesting that the temporal patterns of muscle activation are predominantly defined by body-based cues with possibly only minor reliance on visual cues. N=12.

|  | TAR | GCR | RFR | HLR |
| --- | --- | --- | --- | --- |
| **T<sub>L</sub>V<sub>L</sub>** | 0.87±0.02 | 0.84±0.04 | 0.87±0.02 | 0.86±0.02 |
| **T<sub>L</sub>V<sub>U</sub>** | 0.83±0.02 | 0.82±0.03 | 0.86±0.02 | 0.80±0.04 |
| **T<sub>L</sub>V<sub>L</sub> Increase** | 0.81±0.02 | 0.84±0.03 | 0.80±0.04 | 0.75±0.04 |
| **T<sub>L</sub>V<sub>D</sub>** | 0.83±0.03 | 0.80±0.03 | 0.81±0.04 | 0.86±0.01 |
| **T<sub>L</sub>V<sub>L</sub> Decrease** | 0.70±0.04 | 0.73±0.05 | 0.70±0.04 | 0.64±0.05 |
| **T<sub>U</sub>V<sub>U</sub>** | -0.20±0.09 | -0.17±0.08 | 0.06±0.07 | -0.16±0.07 |
| **T<sub>U</sub>V<sub>D</sub>** | -0.31±0.05 | -0.18±0.06 | 0.05±0.07 | -0.22±0.07 |
| **T<sub>D</sub>V<sub>D</sub>** | 0.61±0.05 | 0.35±0.07 | 0.58±0.06 | 0.53±0.03 |
| **T<sub>D</sub>V<sub>U</sub>** | 0.62±0.04 | 0.33±0.10 | 0.59±0.06 | 0.56±0.05 |

Since we observed changes in muscle activation magnitude in virtual inclinations but not in the patterns, we hypothesized that the source of these magnitude changes is driven by the behavioral gait speed changes seen in the incongruent conditions (Figure 3). Therefore, we evaluated muscle activation magnitude in response to mere changes in gait speed when gravitational cues did not change (see Methods for T_L_V_L_-increase and T_L_V_L_-decrease). Indeed, similar effects on muscle activation magnitudes were seen in the incongruent condition in response to changes in gait speed. For example, in condition T_L_V_D_, the averaged walking speed is decreased (Figure 3) as well as the GCR activation (Figure 6). A similar pattern is illustrated in condition T_L_V_L_-decrease where the speed is decreased in correlation with GCR activation (Figure 7).

**Figure 7.**
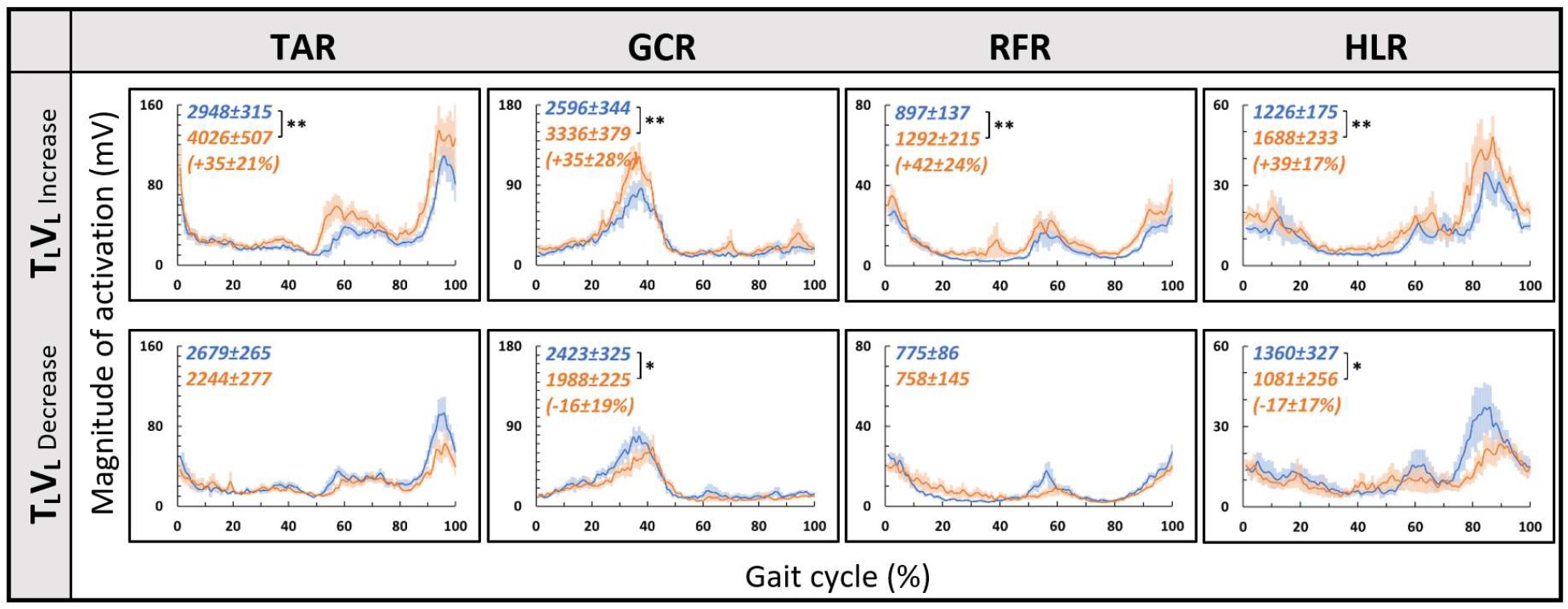
Gait speed does not affect muscle activation patterns for leveled treadmill inclination. Conventions as in Figure 5. Blue lines represent the activity pre-change, and the orange lines represent post-change (increase by ~15% or decrease by ~20% from SSV).

### Visual inclinations do not affect muscle activation patterns

For both the uphill (T_U_, at 10°) and downhill (T_D_, at −10°) treadmill transitions, the parametric manipulation of visual cues did not affect the activation patterns of any of the four muscles (Figure 8A and 8B, respectively). In both treadmill transitions, visual inclination did not make a difference (Table 2). In summary, activation patterns that were modulated by gravitational transitions were insignificantly affected by virtual visual cues.

**Figure 8.**
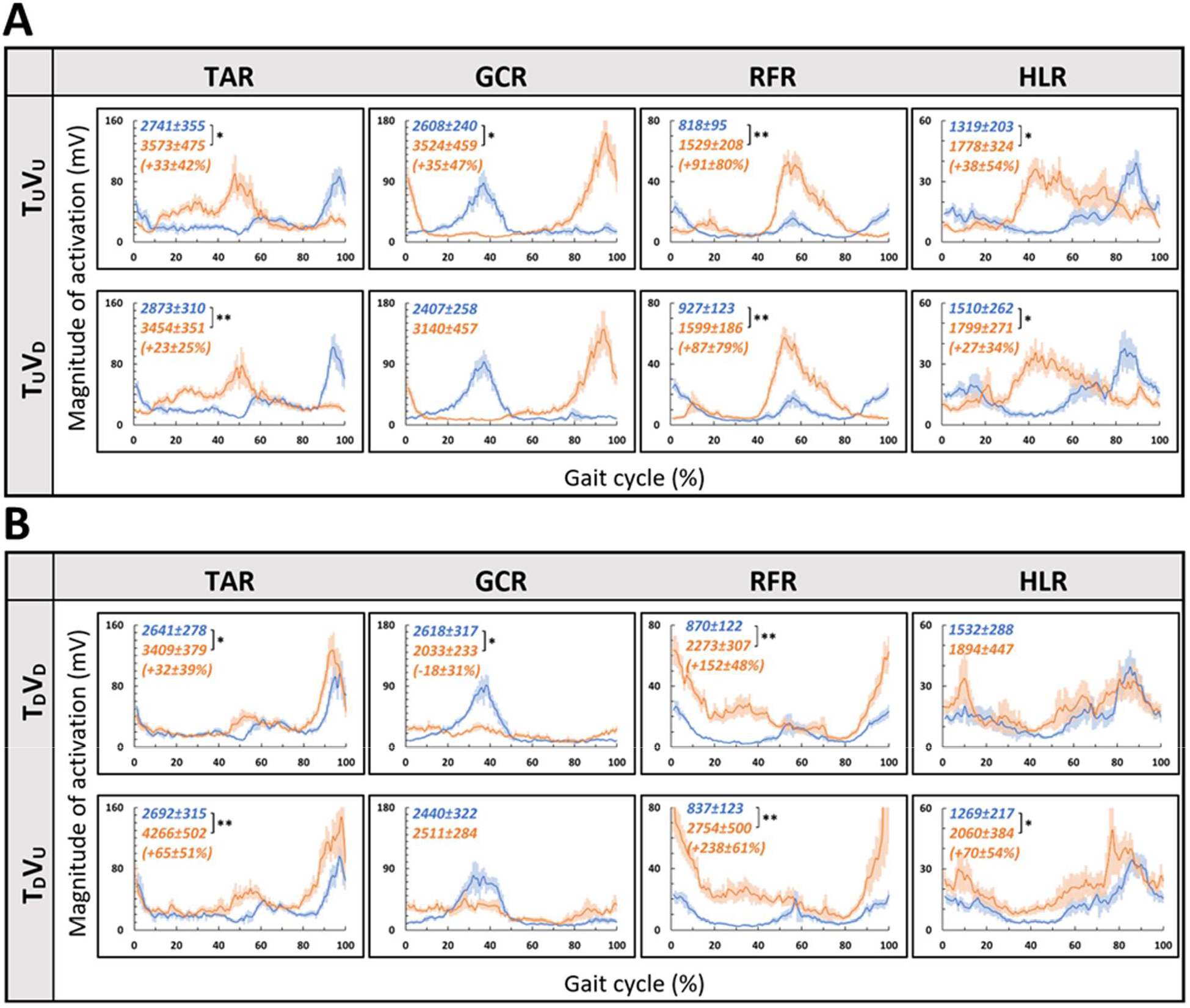
Visual inclinations do not affect muscle activation patterns for uphill (A) or for downhill (B) treadmill inclination. Conventions as in Figure 5. This figure demonstrates that patterns of muscle activation during physical uphill walking did not change regardless of conflicting visual cues (**Panel A:** top row-uphill vision (+10°), bottom row-downhill vision (−10°)) compare orange traces in first and second rows). Likewise, patterns of muscle activation during physical downhill walking were not affected by conflicting visual cues (**Panel B:** top row-downhill vision (−10°), bottom row-uphill vision (+10°) compare orange traces in first and second rows).

By combining the information from Figures 5–8 and Table 2, the observed changes in muscle activation magnitude seen during the incongruent conditions (Figure 6) can be explained by the gait speed changes (Figure 7). This implies that changes in muscle activation temporal patterns are likely to rely predominantly on body-based cues (Figure 8).

## Discussion

In this study, we investigated the contribution of vision to locomotion modulation under gravitational changes. To that end, we manipulated vision independently of body-based cues and measured walking speed and muscle activation. Initially, we replicated recent results that demonstrated that the transition to virtual inclinations modulates locomotion for approximately 20 seconds (Cano Porras et al., 2020), which was evident by speeding up or slowing down (aka the exertion and braking effects, respectively). Furthermore, in an individual differences analysis, we found that the magnitude of this effect was associated with individual visual field dependency. Lastly, we observed that when transitioning from leveled walking to inclined walking, changes in the temporal pattern of muscle activation were primarily affected by gravitational bodily cues (i.e., body-based cues) and not significantly by vision.

According to the internal model of gravity, locomotion modulations regarding gravitational conditions (e.g., walking on inclined surfaces) are based on multiple sensory inputs (visual, vestibular, proprioceptive). However, the proportional contribution of vision under these conditions is unclear. In order to differentiate between the weights of each sensory channel, there is a need to isolate each of them and examine the behavioral and physiological responses. Thus, in real-world conditions, it is difficult to isolate different sensory modalities. One methodology is to recruit specific cohorts as blind people (Dizio and Lackner, 2000), patients with no proprioception (Mullie and Duclos, 2014), or with vestibular impairments (Potter and Silverman, 1984). Another methodology is to artificially manipulate each sensory cue independently, causing a sensory discrepancy. Regarding visual manipulations, most studies change the speed of the projected visual scene in a virtual reality environment, which ultimately affects the participants’ walking speed. These studies show that increasing the optic flow speed in a VR environment causes participants to decrease their walking speed and vice versa (Lamontagne et al., 2007; O’Connor and Donelan, 2012; de Keersmaecker et al., 2019). A recent study examined vision’s contribution to locomotion employing a different visual paradigm: by manipulating physical - visual scene treadmill inclination congruency (Cano Porras et al., 2020). While walking speed was consistently sensitive to visual inputs (supported by our findings as well), it was unclear whether the vision’s contribution extends to affect the magnitude or patterns of muscle activation. Our study found that the magnitude of muscle activation was modulated in relation to the walking speed changes following visually induced simulated virtual inclinations. Such adjustments are in accordance with *indirect prediction* mechanisms, i.e., relying on prior experience to activate pre-programmed gait control patterns that are consistent with expected gravity-based changes (Snaterse et al., 2011; O’Connor and Donelan, 2012; Cano Porras et al., 2020). However, in contrast to changes in gait speed, we found that the pattern of muscle activation during the gait cycle was dominated by body-based cues responding to the actual gravity related (physical) forces but independent of visual cues. This is demonstrated by the observed shift in peak activation during a real gravitational change (Figure 5) but not when the change is only visual (Figure 6). These results are quantitatively corroborated by a high correlation between activation patterns of post- and pre-transition periods (Table 2). Thus, our findings suggest that the vision’s contribution is predominantly involved in dynamic adjustments during indirect prediction for approximately 20 seconds after a perturbation.

While we found that visual cues affect locomotion for a brief period after perturbations occur (i.e., virtual inclination transitions), the role of proprioception and vestibular cues in controlling locomotion are continuous and likely to be dependent on gait speed. Overall, locomotion relies on feedback (e.g., from joint mechanoreceptors that detect gravitational changes) and feedforward (e.g., anticipating center-of-mass change from prior experience)(Riemann and Lephart, 2002; Seidler et al., 2004; Maeda et al., 2019) mechanisms. For example, visual inputs are used in the process of planning locomotion to create a model of the environment in which the walking would occur, while proprioception is essential during the execution of the movement to update the feedforwards commands derived from the visual inputs (Bard et al., 1995; Sainburg et al., 1995). Regarding body-based cues, it seems that during walking, vestibular signals are downregulated (Brandt et al., 1999; Pfaff, 2013; Fabre-Adinolfi et al., 2018), whereas proprioception dominates the pattern of muscle activation (Science and 1985). Overall, neural networks controlling posture and gait are highly dependent on vestibular inputs during slow gait and shift their dependency towards proprioception with increased gait speed. The role of proprioception in these regulations is assumed to be in response to external and internal stimuli (Sorensen et al., 2002). While we found that gait speed was affected by virtual inclination changes, probably due to expectations for gravitational changes (i.e., feedforward effects). Proprioception governed the muscle activation patterns, which were not affected following virtual merely visual inclination manipulations. Internal stimuli are primarily derived by proprioceptive inputs (Tuthill and Azim, 2018), which help adjust the complex interaction between direct muscle activation and indirect muscle activation (movement of one joint inducing movement of another). Again, we demonstrated that before adjusting to a complex mechanism such as walking, the pattern of muscle activation only changes after physical mechanoreceptor-related cue has occurred. We speculate that visual inputs have minimal impact, if any, on neuronal activations that define muscle synergies (Janshen et al., 2017) and temporal muscle activation pattern (Lay et al., 2007; Pickle et al., 2016) characteristics of inclined locomotion. Interestingly, other studies had demonstrated the involvement of visual flow in altering the magnitude of muscle activation during perturbed walking (i.e., when balance control tasks were involved)(Stokes et al., 2017; Cano Porras et al., 2019). For example, Cano Porras et al., recently reported that muscle activation patterns, associated with downward perturbation (walking platforms “drops” down), are also triggered by mere visual perturbations generated in a VR system. In summary, while visual cues have a transiently significant contribution that wears off (O’Connor and Donelan, 2012), body-based cues continuously adjust during walking in response to walking speed.

It is unclear whether visual field dependence is related to or affects locomotion. Patient populations with CNS deficiencies and locomotion deficits show high visual field dependency (measured by subjective visual vertical), probably compensating for their deficits (e.g., Parkinson’s disease (Schindlbeck et al., 2018), post-stroke (Bonan et al., 2006, 2007), multiple sclerosis (Crevits et al., 2007), and cerebral palsy (Slaboda et al., 2013)). Assessments of visual dependency include tests based on visual-vestibular conflicts such as measuring visually-induced illusory perception of self-motion, known as vection (Kim et al., 2015), and the Romberg’s test that measures visually assisted postural stability and aims to identify the influence of vision on postural control by comparing how much the body sways with eyes opened vs. closed (Lê and Kapoula, 2008). In addition to the rod and frame test used in our study, Rothacher et al. (Rothacher et al., 2018) used these two tests and compared them to locomotive outcomes. Out of the three tests, they found that locomotive outcome showed the highest correlation to the rod and frame test by demonstrating that high visual field-dependent participants (as revealed in the rod and frame test) were the most sensitive to visual manipulations in a VR environment. In our study, in line with earlier findings, we found that visual field dependency (as measured by the rod and frame lab-based psychophysical test(Lopez et al., 2006; Isableu et al., 2008; Bagust, 2013) was significantly correlated with locomotive measures. Specifically individuals with higher visual field dependency showed higher percentage of change in gait speed to virtually induced inclinations (e.g., Treadmill up vision down). Together, these findings support the hypothesis that visual field dependency and locomotion outcomes such as visually-dependent gait speed or trajectories are associated and may rely on associated mechanisms.

In this study, we examined the effect of visual cues on gravitational related locomotion modulations in a young, healthy population to explore how muscle activation is affected by visual cues. We found that the magnitude of the behavioral effect (i.e., change in gait speed) was associated with individual visual field dependency. Furthermore, we observed that when transitioning from leveled to inclined walking, the pattern of muscle activation was affected mainly by body-based cues, and not by visual cues. It is still unclear whether the intensity-relations of sensory perceptions of gravity is linear and how people respond to various inclinations, either smaller or larger. A more profound comprehension of locomotion in healthy populations may contribute to understanding pathologies in neurological disorders such as Parkinson’s disease and stroke (commonly presenting with gait disorders). Such knowledge may also be used in combination with VR systems for motor rehabilitation in such conditions (Lamontagne et al., 2007; Darekar et al., 2015; Cano Porras et al., 2018; de Keersmaecker et al., 2019) or in persons with other walking disabilities.

## Acknowledgment

This study was supported in part by the Israel Science Foundation (ISF) grant # 1657-16. SGD was supported by the Israel Science Foundation grant #1485/18. We thank Ms. Shani Kimel-Naor and Mr. Yotam Bahat for their technical assistance.

## Supporting material for the Methods section

### Elaboration on Gait speed-related variables

To assess the post-transition effects on gait speed, we followed the methodologies introduced by Cano-Porras et al. (Cano Porras et al., 2020).

#### Steady-state velocity

A real-time algorithm monitoring treadmill speed determined steady-state velocity (SSV). According to the algorithm, SSV is attained after 1) a minimum 30s of walking, and 2) a consecutive period of 12s with gait speed coefficient of variance less than two percent. Upon satisfying both conditions, the transition of the treadmill and/or visual scene inclination (as appropriate for the experimental condition) was automatically triggered.

#### Normalization of gait speed (Fig. S-1)

Normalization of gait speed (WS) in each experimental condition consisted of three steps. First, WS was divided by the averaged SSV (i.e., from the 12s that defined the SSV period). In the new trace, the mean value of the 12s SSV period is 1. The ratio between WS and the SSV was presented as a percentage. Finally, in order to clearly distinguish between the responses of increased and decreased velocity, the normalized trace was shifted so that the mean value of WS of the SSV period would be zero.

**Figure S-1.**
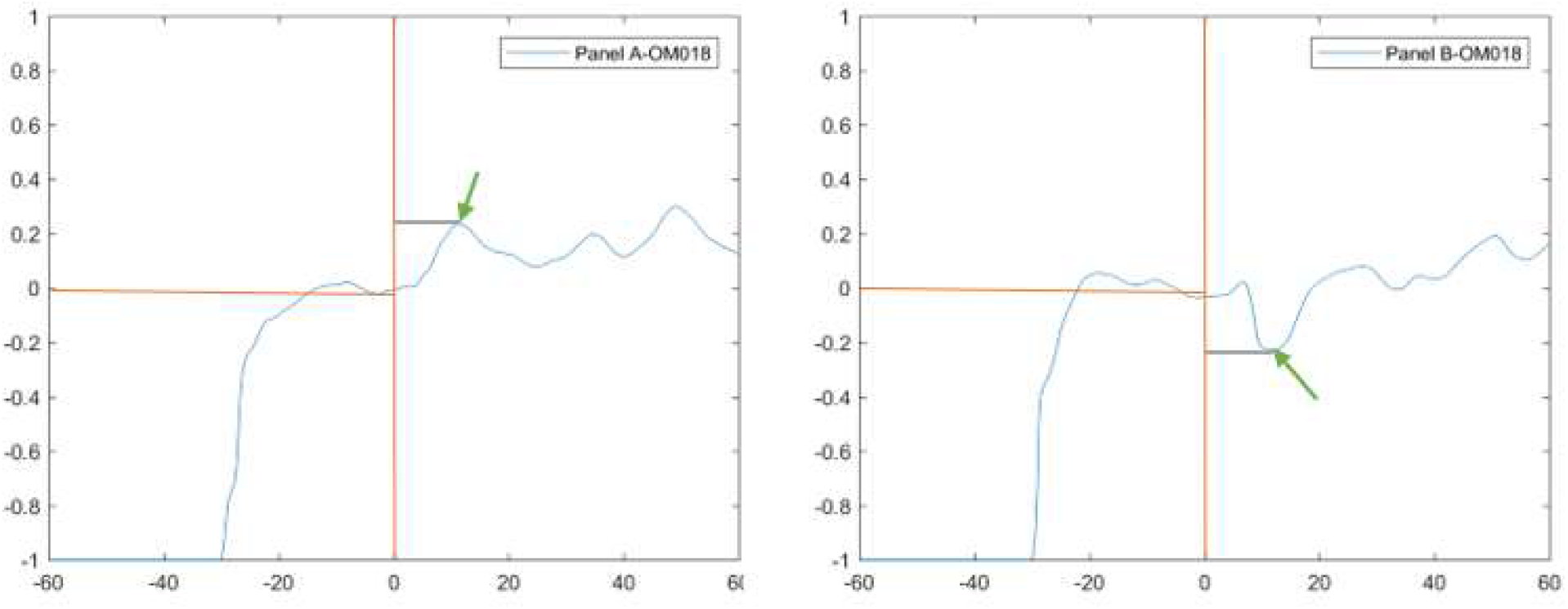
Deriving gait speed-related variables. Fig. S-1 depicts an example of responses to virtual uphill (T_L_V_U_; panel A) and downhill (T_L_V_D_; panel B) transitions from one participant (OM018). The vertical orange line represents the transition, and the horizontal orange line represents steady-state velocity (SSV). On these traces, the peak/trough were identified (green arrows), and the time from transition was calculated (grey line). Units used for magnitude were the relative change (%) in comparison to the SSV values. The second parameter is the time of maximal change (c.f., Fig. S-1).

### Electromyography (EMG) analysis

Data downloaded from the EEGO® system (ANT, The Netherlands) was preprocessed using MATLAB based software programmed in our lab. EMG signals were filtered (finite impulse response band-pass of 20-400 Hz) and full-wave rectified.

To recognize the effect of physical and virtual transitions on muscle activation patterns, we studied the patterns typical to the gait cycle, i.e., the interval between consecutive heel strikes. Therefore, gait cycle timing was first identified from the ground reaction force signals obtained by the force plates embedded beneath the treadmill’s belts. We identified the gait cycles 10 seconds before the transition (when the participant walked in SSV), during the 5 seconds of transition and 18 seconds post-transition (Figure S-2). We then chose by visual inspection 5-9 cycles for the pre and post-transition (purple and orange frames in the figure). The criterion of choice was to avoid residual effects of the 5 seconds of transition (green frame in the figure). According to the example of T_U_V_U_ condition, the EMG pattern through all the frames is roughly similar before the transition, compared to a gradual change of EMG pattern during the transition. It is critical to note that after the transition, the EMG pattern is constant throughout the selected gait cycles.

We compared pre-transition to post-transition EMG patterns for each subject. For this, all selected gait cycle times were re-scaled (0-100% of the gait cycle, between two consecutive heel strikes). From the time-scaled EMG traces, we grand averaged all the pre-data and post-data separately (see Fig. S-3). With the averaged EMG traces, we computed the following two parameters:

#### (1) Magnitude

the summation (i.e., area under the curve) of the EMG traces. In this example, the MAG values are depicted in the upper left side of the panel (Fig S3). Also, the normalized percentage of the difference is stated in parenthesis, where a significant change was observed between post- and pre-transitions.

### Normalization for summation of the magnitude of activation

Multiple participants in the study may require different scaling due to different conditions (e.g., sweating, hair, conductivity). To address this problem, each participants’ results were normalized per condition according to the value of the magnitude of activity in the pre-transition condition.

#### (2) Similarity of EMG activation pattern

The post transition EMG trace was correlated with the pre-transition EMG trace (Pearson Correlation) for each participant for each condition. A high correlation value indicates a small change in the pattern of activation. In this example (Fig S3), the correlation value is −0.17±0.08 between the orange and blue trace. This low value indicates a significant change in EMG activation pattern seen between leveled walking and uphill walking in the gastrocnemius (GCR) muscle.

**Figure S-2.**
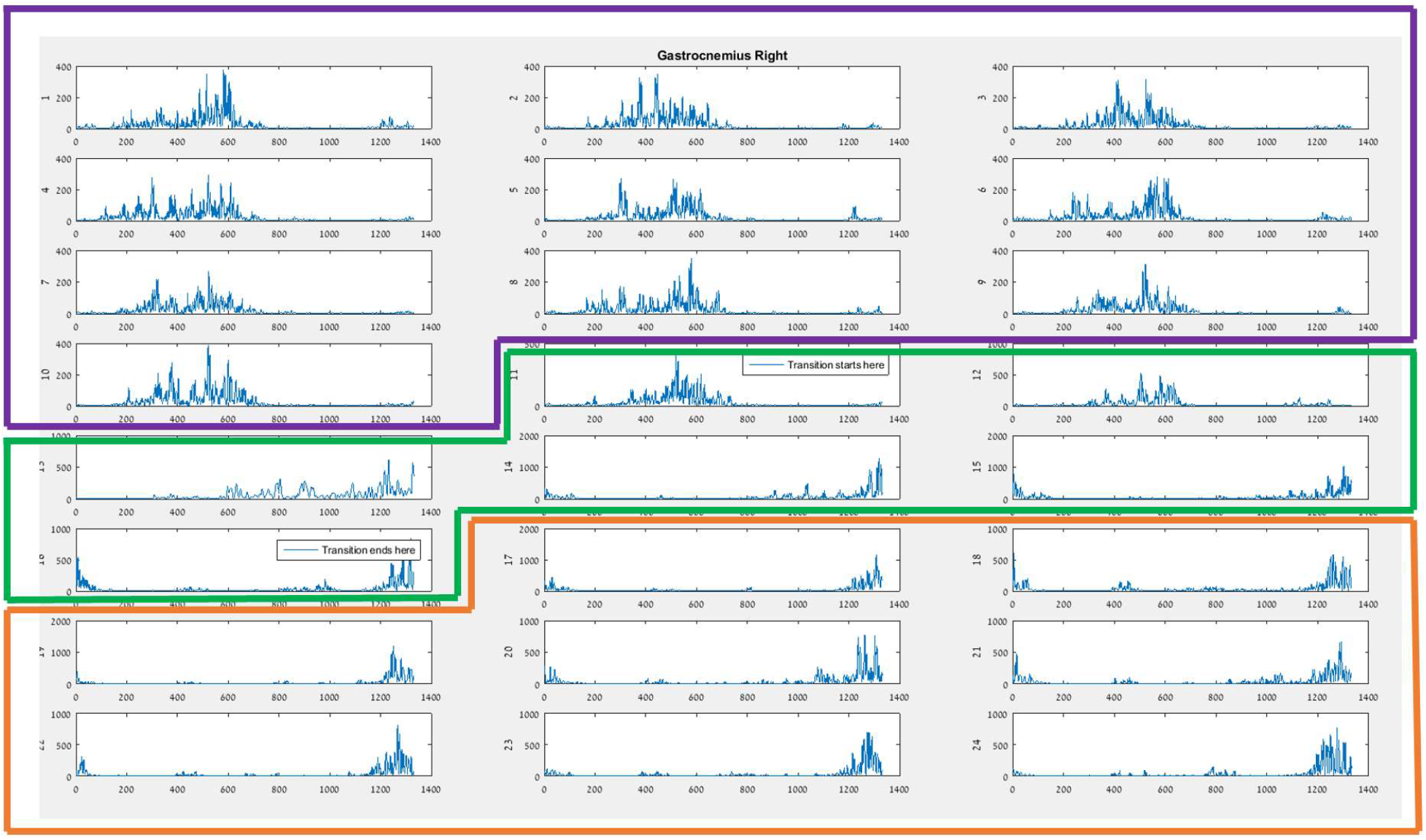
A graphical representation of the right gastrocnemius (GCR) muscle from 24 gait cycles obtained from one participant during the T_U_V_U_ condition. The X-axis shows the sample rate [Hz], and the Y-axis shows muscle activation [mV]. As can be seen, the pattern of activation remains similar for all the gait cycles pre-transition (purple frame) and post-transition (orange frame); the green frame represents the 5s transition.

**Figure S-3.**
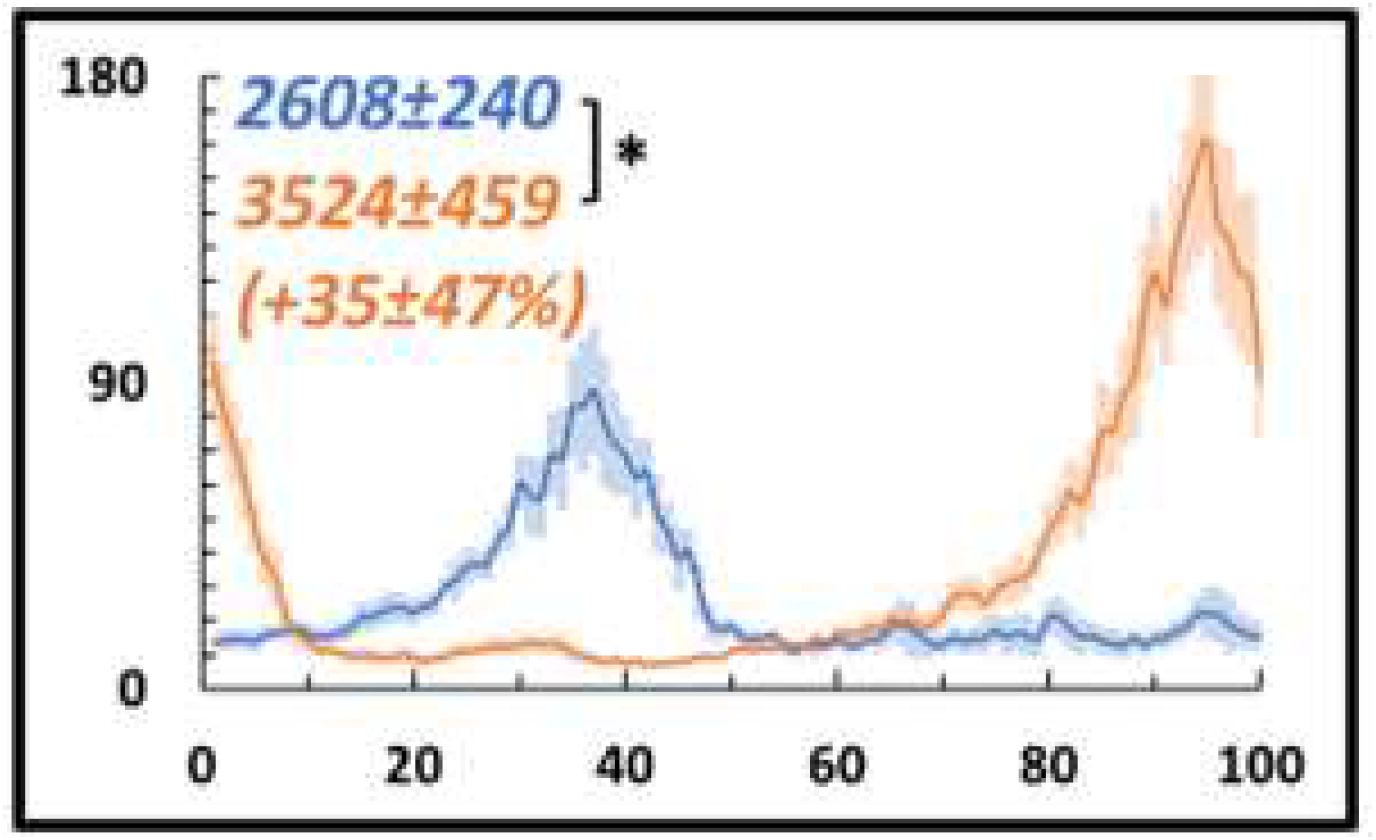
Pre- vs. post- EMG right Gastrocnemius activation patterns for visual-physical congruent uphill walking. Grand average (across participants, N=12) of muscle activation patterns (Y-axis, in mV) plotted against a time-normalized gait cycle on the X-axis (in % of the gait cycle) in blue (shaded areas represent SE) for pre-transition and in orange (shaded areas represent SE) for 20 seconds post-transition. Numerical values at top left represent averaged summation (area under the curve) magnitude of activation ± SE for pre-transition (blue) and post-transition (orange), asterisks denote a significant post- vs. pre-transition change (two-tailed, paired T-test, P<0.05=*). The normalized change appears below (in %) in parentheses.

## Supporting material for the Results section

### Reproducing previous results

Table S-1 provides comparison between the results of the present study and the results of an earlier study conducted in this laboratory using the same paradigm. The following data confirms the reproducibility of results using the present virtual reality paradigm.

**Table S-1.** No significant difference in the patterns of muscle activation between Cano Porras 2020 and the recent study. Each row represents a walking condition (see methods for details). The 2^nd^ and 3^rd^ columns represent the average values for the relative peak change from SSV (%)±SE. The 5^th^ and 6^th^ columns represent the average time of peak effect (seconds) for each study±SE. The 4^th^ and last columns represent the p-value obtained from a paired two-tailed t-test. * (N=14), ** (N=12).

|  | Magnitude of relative speed change (%) |  |  | Time of maximal effect (s) |  |  |
| --- | --- | --- | --- | --- | --- | --- |
|  | Cano Porras<br>2020* | Present<br>study** | P-value | Cano Porras<br>2020* | Present<br>study** | P-value |
| T<sub>U</sub>V<sub>U</sub> | 32.8±15.4 | 25.4±12 | 0.18 | 14.4±6.8 | 13.5±5.1 | 0.69 |
| T<sub>D</sub>V<sub>D</sub> | 13.9±23.3 | 18.8±14.1 | 0.47 | 17.2±8.2 | 22.2±8.3 | 0.15 |
| T<sub>L</sub>V<sub>U</sub> | 15.9±11.2 | 17.9±8.9 | 0.64 | 12.3±4.5 | 10.3±2.2 | 0.19 |
| T<sub>L</sub>V<sub>D</sub> | 20.7±16.8 | 25±15.5 | 0.5 | 7.6±1.9 | 8±2 | 0.59 |
| T<sub>D</sub>V<sub>U</sub> | 21.6±17.2 | 28.1±20 | 0.38 | 11.6±5.5 | 9.2±2.7 | 0.16 |
| T<sub>U</sub>V<sub>D</sub> | 62.8±24.7 | 58.3±30.4 | 0.68 | 7.9±1.7 | 7.2±2.3 | 0.38 |

#### Supplementary Video 1

https://drive.google.com/file/d/1FLQsbuMKCX7xKWtewG89PfDTI7WSM19I/view?usp=sharing

The video insert depicts the behavior effects of visually induced downhill walking. Initially, the person is walking with the treadmill, and the visual scene leveled. Once the person reaches steady-state velocity and maintains it for 12 seconds, a downward transition of the visual scene occurs, while the treadmill remains leveled. Note how she decreases her gait speed to counteract the expected gravitational forces (i.e., the braking effect). Eventually, body-based cues govern, and gait speed returns to the previous steady-state values.

